# Assessment of the Impact of Sleep Deprivation and Diet On Growth, Cognition, Inflammation, Oxidative Stress, and Tissue Histology

**DOI:** 10.1101/2024.10.04.616642

**Authors:** Lutfat A. Usman, Emmanuel O. Ajani, Afolabi C. Akinmoladun, Rasheed B. Ibrahim, Bukola R. Oyedeji, Toheeb A. Adedotun, Ganiyat A. Mustapha, Ramat Y. Abduwahab, Awanat I. Banuso, Oladapo O. Oyedokun, Ayobami R. Olatunji, Kolawale Akande, Opeyemi Grace Oni, Olamide Oduola, Oladimeji B. Oguntayo, Dauda K. Saka, Abdul-Hafeez O. Hussain

## Abstract

This study explores the interplay encompassing amino acids’ roles, metabolic processes, growth, and the intricate relationships between sleep patterns, amino acid-deficient diet, and caffeine consumption, with emphasis on the vital connections between diet, oxidative stress, and cognitive health while investigating the essential role of quality sleep in maintaining cognitive function. This study investigates the intricate connections among amino acids, metabolic processes, growth, sleep patterns, amino acid-deficient diets, and caffeine consumption. Emphasizing the crucial links between diet, oxidative stress, and cognitive health, the research underscores the essential role of quality sleep in maintaining cognitive function. The bidirectional relationship between disrupted sleep and cognitive health, as well as the impact of caffeine on cognitive function, is explored. The study involved adult male Wistar rats divided into 10 groups A - J based on cage conditions and diet. Group A-E were kept in a normal cage with diets containing varying levels of tryptophan and other essential amino acids, while Group F-J underwent sleep deprivation using a disk-over-water method and received diets with varying tryptophan and other essential amino acid levels, in addition to caffeine. The experiment spanned 2 weeks, blood samples were collected on days 1,4,7, and 13 for relevant biochemical serum analyses, and the rats were sacrificed on the fourteenth day for blood sample collection, biochemical assays, and brain histology. The 2-week experiment revealed a linear decrease in melatonin and serotonin levels across all caffeine-administered groups, possibly due to stress and anxiety as compared with the reduction in these parameters observed in control rats, suggesting melatonin and serotonin as potential sleep deprivation biomarkers. Testosterone and insulin-like growth hormone assays showed a significant decrease in sleep-deprived rats with caffeine, impacting secondary sex characteristics and growth. Elevated TNF-α and interleukin-1b levels in the caffeine-administered, sleep-deprived group suggest potential health risks associated with sleep deprivation and caffeine. Serum assays for antioxidant enzymes indicated heightened oxidative stress in sleep-deprived rats with caffeine. Histopathological studies revealed cellular abnormalities in the sleep-deprived group, indicating increased cellular metabolism and potential pathological conditions. This study elucidates the complex relationships among sleep, dietary factors, and hormonal and inflammatory responses. It underscores the potential neurotoxic effects associated with caffeine consumption when coupled with sleep deprivation. These results underscore the urgent need for further investigation into dietary strategies that could alleviate these adverse effects and promote better health.

## INTRODUCTION

In today’s fast-paced society, many individuals rely on caffeine to maintain wakefulness and alertness, often accompanied by a nutritionally deficient diet consumed while on the move. This routine, characterized by insufficient rest and poor nutrition, perpetuates a cycle of chronic fatigue and suboptimal health. Amino acids, essential for protein synthesis and the production of neurotransmitters and hormones, play a critical role in maintaining physiological functions. Of the 20 amino acids, nine are considered essential, including phenylalanine, valine, tryptophan, threonine, isoleucine, methionine, histidine, leucine, and lysine. These essential amino acids must be obtained through diet, as the body cannot synthesize them. In specific conditions, such as pregnancy, adolescent growth, or recovery from trauma, other amino acids like arginine and histidine become conditionally essential due to increased physiological demands (Moro et al., 2020). Protein quality is measured by its ability to meet the nutritional requirements of essential amino acids (EAAs). Deficiencies in EAAs, such as lysine and threonine, adversely affect growth, bone development, and protein metabolism, as demonstrated in animal models (Moro et al., 2020).

The adult human brain, which constitutes only 2% of body weight, requires approximately 20% of the body’s energy, primarily derived from glucose, to sustain neurotransmission. Unlike skeletal muscles, which store glycogen, the brain relies on a constant supply of glucose from the bloodstream, as it has minimal energy reserves. In conditions of reduced glucose availability, the brain can utilize alternative substrates such as lactate, medium-chain triglycerides, and ketone bodies. The brain’s high metabolic activity is maintained even during sleep, with local neuronal activity varying significantly depending on sensory or motor stimulation and wakefulness transitions. The phenomenon of ‘neurometabolic coupling’ ensures that local energy supply and cerebral blood flow are adjusted according to neuronal activity, thereby maintaining proper brain function (Tardy et al., 2020).

Diet and mental health are closely linked, with evidence suggesting that amino acids such as tryptophan play a critical role in mood regulation and cognitive function. Tryptophan, a precursor to serotonin (5-HT), is found in protein-based foods and is protective against depression. Severe dietary restrictions, as seen in anorexia nervosa, are associated with depressive symptoms and impaired cognitive functions, highlighting the importance of adequate nutrition for mental health. Additionally, tryptophan depletion has been shown to affect cognitive tasks such as response times and visual discrimination, further emphasizing its role in cognitive health(Reuter et al., 2021).

The impact of caffeine on cognitive function is complex and multifaceted. While caffeine has been shown to have short-term benefits in improving memory and cognition, its long-term effects are less clear, with studies presenting conflicting results regarding its association with cognitive decline and dementia risk (Domínguez et al., 2021). Sleep deprivation, often exacerbated by caffeine consumption, has been linked to impaired immune function, increased cancer risk, and disrupted cognitive processes. Sleep is essential for overall health, with non-rapid eye movement (NREM) and rapid eye movement (REM) stages serving distinct functions, including memory consolidation and growth hormone release (Mason et al., 2021). Sleep deprivation is also associated with memory deficits, possibly due to increased oxidative stress in the hippocampus, and a Western diet, high in fats and sugars, exacerbates these effects by increasing oxidative stress and lipid peroxidation (Mason et al., 2021).

In this study, we investigated the hypothesis that a combination of sleep deprivation and tryptophan-deficient diet exacerbate the learning and memory impairment induced by each factor alone. Molecular approaches using enzymatic and colorimetric assays were used to test this hypothesis.

## METHODOLOGY

## EXPERIMENTAL FEED PROCUREMENT AND PROCESSING

### Experimental Feed Preparation and Analysis

The experimental feeds utilized in this study included standard animal chow (Topfeed), milled maize, and pelletized red and white sorghum. These feeds were sourced from the Malete farm market, while the maize and sorghum were manually cleaned to remove any visible farm dirt. Subsequently, the cleaned maize and sorghum were milled and pelletized at an Animal Feed Mill, located in Ilorin, Kwara State. Each feed type underwent proximate and amino acid composition analysis to evaluate and compare their nutritional profiles. The analyses were carried out in triplicates (n=3).

### Determination of Amino Acid Profile

The amino acid profile of the experimental feeds was determined following the method described by AOAC (2005) with slight modifications. The sample was dried at 70°C to constant weight, defatted, hydrolyzed, evaporated using a rotary evaporator, and then analyzed with an Applied Biosystems PTH Amino Acid Analyzer.

### Defatting the Sample

Approximately 4 g of the sample was defatted using a chloroform/methanol mixture in a 2:1 ratio. The sample was placed in an extraction thimble or wrapped in filter paper and subjected to 15 hours of extraction using a Soxhlet apparatus, as described by AOAC (2006).

### Nitrogen Determination by Micro Kjeldahl Method

Nitrogen content was determined using the micro Kjeldahl method, with slight modifications from the AOAC (2005) protocol. In this process, nitrogen from proteins and other compounds is converted to ammonium sulfate by acid digestion with boiling sulfuric acid. 250 mg of the sample was placed in a Kjeldahl flask, and 200 mg of a catalyst mixture (potassium sulfate, copper sulfate, and selenium powder) was added. After adding 10.0 cm³ of concentrated sulfuric acid, the mixture was gently heated until frothing ceased, then further heated for 1 hour and 30 minutes. The digest was cooled and diluted to 100 cm³ with distilled water.

A 10.0 cm³ aliquot of the diluted digest was distilled in a micro Kjeldahl distillation apparatus with the addition of 10.0 cm³ of 40% sodium hydroxide solution. The distillate was collected in 10.0 cm³ of 4% boric acid containing a mixed indicator. The color change from red to green was noted, and the solution was titrated with standard 0.01N or 0.02N hydrochloric acid to a grey endpoint. The nitrogen percentage was calculated using the following formula:

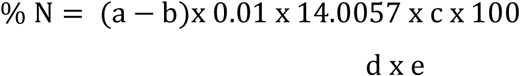

a = Titre value for the sample

b = Titre value for the blank

c = Volume to which digest is made up with distilled water

d = Aliquot taken for distillation

e = Weight of dried sample (mg)

### Hydrolysis of the Sample

250 mg of the defatted sample was placed into a glass ampoule, and 7 ml of 6N HCl was added. Oxygen was expelled by passing nitrogen into the ampoule to prevent oxidation of sensitive amino acids like methionine and cystine during hydrolysis. The ampoule was sealed with a Bunsen burner flame and heated in an oven at 105°C ± 5°C for 22 hours. After cooling, the ampoule was carefully opened, and the contents were filtered to remove humins. (Note: Tryptophan is destroyed by 6N HCl during hydrolysis.) The filtrate was evaporated to dryness using a rotary evaporator, and the residue was dissolved in 5 ml of acetate buffer (pH 2.0).

### Loading the Hydrolysate into the Analyzer

A 60-microliter aliquot of the hydrolysate was dispensed into the cartridge of the analyzer. The Applied Biosystems PTH Amino Acid Analyzer, designed to separate and analyze free acidic, neutral, and basic amino acids, was used to determine the concentration of each amino acid in the hydrolysate. The period of analysis lasted for 45 minutes.

### Calculation of Amino Acid Values

An integrator attached to the analyzer calculates the peak area, which is proportional to the concentration of each amino acid present in the sample.

### Tryptophan Hydrolysis and Determination

For the determination of tryptophan, the sample was hydrolyzed using 4.2 M NaOH, following the method by Maria et al. (2004). The sample was first dried to a constant weight and defatted using a chloroform/methanol mixture (2:1 ratio) in a Soxhlet extraction apparatus for 15 hours, as described by AOAC (2006). After defatting, 250 mg of the sample was weighed into a glass ampoule. To this, 10 mL of 4.2 M NaOH was added, and oxygen was expelled by passing nitrogen into the ampoule to prevent oxidation of amino acids during hydrolysis. The ampoule was then sealed with a Bunsen burner flame and placed in an oven preset at 105°C ± 5°C for 4 hours. Following hydrolysis, the ampoule was cooled, opened at the tip, and its contents filtered to remove humins. The filtrate was then neutralized to pH 7.0, evaporated to dryness under vacuum at 40°C using a rotary evaporator, and the residue dissolved in 5 mL of borate buffer (pH 9.0). The hydrolysate was prepared for analysis by loading 60 µL into the cartridge of the analyzer. An integrator attached to the analyzer calculated the peak area, which is proportional to the concentration of tryptophan.

## PROXIMATE ANALYSIS OF MOISTURE, PROTEIN, FAT, ASH, AND CRUDE FIBRE

### Moisture Content Determination (AOAC, 2006, Modified)

The sample was thoroughly mixed before analysis. Approximately 2 g of the sample was weighed into a glass petri dish, which had been pre-dried and weighed. The dish containing the sample was then placed in a hot air oven at 130°C ± 3°C for 5 hours. The sample was dried to a constant weight, cooling for ten minutes in a desiccator before each weighing.

The percentage of moisture was calculated using the following formula:

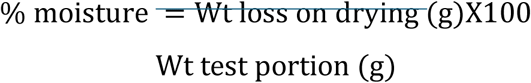

Where:

- Wt = Weight of the sample (g)

### Nitrogen Determination by Micro Kjeldahl Method (Crude Protein) (AOAC, 2005)

The nitrogen in proteins and other compounds was converted to ammonium sulfate through acid digestion with boiling sulfuric acid. A known weight (200 mg) of the sample was placed in a Kjeldahl flask, and approximately 200 mg of a catalyst mixture (potassium sulfate, copper sulfate, and selenium powder) was added. 10 cm³ of concentrated sulfuric acid was added to the flask. The mixture was heated gently until frothing ceased and then further digested for 1 hour. After cooling, the digest was diluted to a known volume (100 cm³) with distilled water.

A 10.0 cm³ aliquot of the diluted digest solution was distilled using a micro Kjeldahl distillation apparatus, with 10.0 cm³ of 40% sodium hydroxide solution added for steam distillation. The distillate was collected in 10.0 cm³ of 4% boric acid containing a mixed indicator and titrated with standard 0.01N or 0.02N hydrochloric acid to a grey endpoint.

The percentage of nitrogen was calculated using the following formula:

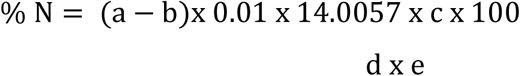

a = Titre value for the sample

b = Titre value for the blank

c = Volume to which digest is made up with distilled water

d = Aliquot taken for distillation

e = Weight of dried sample (mg)

To convert to % crude protein, the nitrogen percentage is multiplied by the conversion factor (6.25).

### Ash Determination (AOAC, 2005, Method 942.05)

Two grams (2 g) of the test portion was weighed into a porcelain crucible and placed in a muffle furnace preheated to 600°C. The sample was incinerated at this temperature for 2 hours. After incineration, the crucible was transferred directly to a desiccator, cooled, and weighed immediately. The percentage of ash was reported to two decimal places.

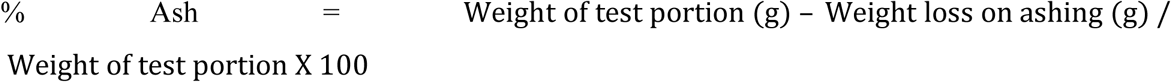

### Fat Determination (Ether-Extract) (AOAC, 2005, Method 945.16, Modified)

A Soxhlet extraction apparatus and a 250 ml quick-fit flask, pre-dried in an oven, were used for fat extraction. Approximately 5 g of the sample was weighed and transferred to a fat-free extraction thimble, lightly plugged with cotton wool. The thimble was placed in the extractor, and 150 cm³ of petroleum ether (boiling point 40-60°C) was added to the flask. The extraction was carried out for at least 6 hours with gentle boiling using an electrothermal heating mantle.

After extraction, the ether was evaporated at 100°C, and the residue was dried in an air oven for 1 hour at 100-105°C. The percentage of fat was reported to the second decimal place.

% Fat = Weight of Fat Extract (g)/Weight of Sample (g) X 100

### Crude Fibre Determination (AOAC, 2005, Method 978.10, Modified)

The defatted ground sample from the fat determination was transferred into a 250 ml quick-fit flask. Then, 150 ml of 1.25% sulfuric acid was added, and the mixture was refluxed for 30 minutes. After cooling, the mixture was filtered using a Buchner funnel fitted with Whatman filter paper and rinsed three times with hot distilled water. The residue was transferred into the flask, and 150 ml of 1.25% sodium hydroxide was added, followed by refluxing for another 30 minutes.

The mixture was filtered again using a Buchner funnel, rinsed with hot distilled water, followed by 1.25% sulfuric acid, and finally with 95% ethanol. The filter paper containing the residue was placed in a porcelain crucible, dried in an oven for 2 hours at 130°C, cooled in a desiccator, ashed at 550°C ± 10°C in a muffle furnace, cooled again in a desiccator, and weighed. The crude fibre content was calculated and reported as a percentage.

% Crude Fiber = (Weight of Residue after digestion (g) − Weight of Ash (g)/ Weight of Sample (g))X 100

## EXPERIMENTAL DESIGN

The animal protocol used in this study was approved by the Faculty of Pure and Applied Sciences Animal Ethics Committee, following recommendations from the Research Committee at Kwara State University, Malete. Eighty adult male Wistar rats (weighing 145g-150g) were obtained from the animal holding unit of the Biochemistry Department at Summit University, Kwara State, Nigeria, and acclimatized for one week. The rats were then divided into two sets, labeled groups A-J, with eight rats in each group as outlined in Table 1. The experimental phase, during which the dietary and housing conditions were implemented according to the table, was conducted for two weeks. Caffeine powder was purchased from First Octopus Chemicals, Ogbomosho, Nigeria. All other reagents used were of analytical grade, and the water used in the experiments was glass distilled.

**Table 1:** Summary of Experimental Group Treatments Based on Dietary and Housing Condition Variations.

| Group | Housing Condition | Diet | Caffeine Administration |
| --- | --- | --- | --- |
| A | Regular animal cage housing | Regular animal chow and water ad libitum | None |
| B | Regular animal cage housing | Milled maize and water ad libitum | None |
| C | Regular animal cage housing | Pelletized sorghum and water ad libitum | None |
| D | Regular animal cage housing | Milled maize and water ad libitum | Bi-daily administration of 20 mg/kg body weight of caffeine powder |
| E | Regular animal cage housing | Pelletized sorghum and water ad libitum | Bi-daily administration of 20 mg/kg body weight of caffeine powder |
| F | Disk-over-water cage housing | Regular animal chow and water ad libitum | None |
| G | Disk-over-water cage housing | Milled maize and water ad libitum | None |
| H | Disk-over-water cage housing | Pelletized sorghum and water ad libitum | None |
| I | Disk-over-water cage housing | Milled maize and water ad libitum | Bi-daily administration of 20 mg/kg body weight of caffeine powder |
| J | Disk-over-water cage housing | Pelletized sorghum and water ad libitum | Bi-daily administration of 20 mg/kg body weight of caffeine powder |

## SLEEP DEPRIVATION PROCEDURE

The disk-over-water method was utilized to induce sleep deprivation while minimizing the effects of social isolation by housing two rats each, in separate compartments sharing a common round disk that formed an elevated partial floor, with shallow water covering the cage floor beneath the disk. The cages were constructed from glass, with punctured holes at the top to allow for respiratory exchange. Care was taken not to puncture the bottom of the cages to prevent the rats from using the holes to climb up and escape the rotating platform. The cages were constructed from glass, with punctured holes at the top to allow for respiratory exchange. Care was taken to avoid puncturing the bottom of the cages, preventing the rats from using the holes to climb up and escape the rotating platform.

The disk automatically rotated at a slow speed in response to the rats’ movements. When a sleep-deprived rat fell asleep and lost muscle tone, it would fall into the water, triggering wakefulness and disturbing the balance of the rotating disk, which could cause the other rats to also fall into the water. The rats would then swim out of the water and return to the platform. A 2-hour daily break was provided, during which a small, non-rotating platform was placed beside the rotating disk, allowing the rats to sleep without falling into the water. The rats were pre-trained for one week before the experiment began to recognize the non-rotating platform as a steady platform. After the 2-hour break, the non-rotating platform was removed, and the sleep deprivation procedure resumed. This protocol was maintained for two weeks, resulting in the sleep-deprived rats achieving only approximately 16% of their normal sleep.

## BLOOD SAMPLE PREPARATION AND ANIMAL SACRIFICE AND BLOOD SAMPLE PREPARATION

Blood samples were collected from rats via tail vein in each group on days 1, 4, 7, 10, and 13 to carry out biochemical assays. Immediately after the two-week experimentation period, the animals were anesthetized with diethyl ether. The animals were dissected immediately, and the brain was carefully excised for histomorphology analyses.

## ELISA ASSAY FOR TESTOSTERONE, IGF-1, SEROTONIN, MELATONIN, INTERLEUKIN-1 Β, AND TNF-Α

The concentrations of testosterone, IGF-1, serotonin, melatonin, interleukin-1 beta, and TNF-alpha were determined in the prepared plasma using enzyme-linked immunosorbent assay (ELISA) kits, employing the sandwich ELISA principle.

### Principle

In this assay, each well on the microtiter plate had been pre-coated with a specific antibody. Standards or samples were added to the wells, allowing the target antigen to bind to the antibody. Unbound substances were subsequently washed away. A biotin-conjugated detection antibody, specific to the captured antigen, was then introduced. After washing to remove any unbound detection antibody, an avidin-horseradish peroxidase (HRP) conjugate was added, which bound to the biotin. A tetramethylbenzidine (TMB) substrate was introduced, leading to color development through its reaction with the HRP enzyme. The reaction was halted by the addition of a sulfuric acid stop solution, which caused a color change from blue to yellow. The optical density (OD) of the wells was measured at a wavelength of 450 nm. The OD values were then compared to a standard curve generated with known antigen concentrations to determine the antigen levels in the samples.

### Procedure

1. Testosterone and IGF-1 Assays:

1. 50 μL of standard or sample was added to each well.
2. 50 μL of HRP conjugate was then added, and the mixture was incubated for 1 hour at 37°C for the testosterone assay, and 2 hours at 37°C for the IGF-1 assay.
3. The wells were washed three times with 250 μL of wash buffer.
4. 50 μL of substrate A and substrate B working solutions were added to each well, and the plates were incubated for 15 minutes at 37°C, protected from light.
5. Finally, 50 μL of stop solution was added, and the OD was measured at 450 nm.
2. Melatonin and Serotonin Assays:

1. 50 μL of the sample was added to each well.
2. For the melatonin assay, 50 μL of HRP conjugate was added, followed by incubation for 1 hour at 37°C, after which the wells were washed twice.
3. For the serotonin assay, 100 μL of HRP conjugate was added, followed by incubation for 1 hour at 37°C, with the wells then washed four times.
4. 50 μL of chromogen solution A and 50 μL of chromogen solution B were added to each well, followed by incubation for 15 minutes at 37°C, protected from light.
5. 50 μL of stop solution was then added, and the OD was measured at 450 nm.
3. Interleukin-1 β Assay:

1. 100 μL of standard, blank, or sample was added to each well, and the plate was incubated for 2 hours at 37°C.
2. The liquid was aspirated without washing, and 100 μL of Detection Reagent A working solution was added, followed by incubation for 1 hour at 37°C.
3. The wells were washed three times with 350 μL of wash buffer.
4. 100 μL of Detection Reagent B working solution was added, and the plate was incubated for 30-60 minutes at 37°C, followed by five washes.
5. 90 μL of TMB substrate solution was then added, and the plate was incubated for 15-30 minutes at 37°C, protected from light.
6. 50 μL of stop solution was added, and the OD was measured at 450 nm.
4. TNF-α Assay:

1. 100 μL of standard or sample was added to each well, and the plate was incubated for 90 minutes at 37°C.
2. The liquid was removed, and 100 μL of biotinylated detection antibody working solution was added, followed by incubation for 1 hour at 37°C.
3. The wells were washed three times with 350 μL of wash buffer.
4. 100 μL of HRP conjugate working solution was added, and the plate was incubated for 30 minutes at 37°C, followed by five washes.
5. 90 μL of TMB substrate solution was added, and the plate was incubated for 15 minutes at 37°C, protected from light.
6. 50 μL of stop solution was added, and the OD was measured at 450 nm.

## EVALUATION OF OXIDATIVE STRESS

The evaluation of oxidative stress in the serum of the Wistar rats in each experimental group was determined using the method described by Munteanu and Apetrei, 2021with modifications.

### Determination of Superoxide Dismutase (SOD) Activity

SOD accelerates the dismutation of superoxide radicals (O2-), formed during oxidative energy production, into hydrogen peroxide and molecular oxygen. This method is based on the measurement of optical density resulting from the reaction of xanthine and xanthine oxidase, where superoxide radicals react with nitroblue tetrazolium (N.B.T) to form a blue-colored formazan dye. The optical density was read at a wavelength of 560 nm. The SOD present in the serum sample inhibits the formation of formazan by scavenging superoxide radicals from the environment.
*SOD Activity (U/L) = (ΔA/min)/ (ε × d)*
Where: ΔA = change in absorbance at 560 nm, ε = molar absorptivity of the product formed and d = path length of the cuvette in cm

### Determination of Catalase (CAT) Activity

The concentration of the catalase enzyme was determined by measuring the absorbance of a solution containing 1.4 mL of 30% hydrogen peroxide (H2O2), 0.1 mL of phosphate buffer, and 1.4 mL of the sample. The absorbance was measured every 30 seconds at a wavelength of 240 nm. The enzyme activity in units per liter (U/L) was calculated using the following formula:

CAT Activity (U/L) = (2.3 / Dx) × (log A1 / log A2)

Where Dx represents the time interval of 30 seconds, A1 is the initial absorbance, and A2 is the final absorbance.

### Determination of Reduced Glutathione (GSH) Concentration

The concentration of reduced glutathione (GSH) was measured in EDTA-treated blood after hemolysis with distilled water. Proteins lacking sulfhydryl (SH) groups were precipitated using a precipitation solution. The GSH content was then determined based on the formation of yellow color in the reaction between sulfhydryl groups and 5,5’-dithiobis-(2-nitrobenzoic acid) (DTNB). The optical density of the reaction was measured at a wavelength of 412 nm using a spectrophotometer within 24 hours. The GSH activity was calculated using the following formula:

Concentration (mg/dL) = [(OD2 – OD1) / (13600 × E1)] × 1.25 × 1000

Where OD1 is the first absorbance measured before the addition of DTNB at 412 nm, OD2 is the second absorbance after the addition of DTNB at 412 nm, and 13600 is the molar extinction coefficient of the yellow color formed during the reaction between GSH and DTNB.

### Determination of Malondialdehyde (MDA) Concentration

The concentration of malondialdehyde (MDA), a final product of lipid peroxidation, was determined by measuring its reaction with thiobarbituric acid (TBA), which forms a colored complex. A 200 µL blood sample was mixed with 800 µL phosphate buffer, 25 µL butylated hydroxytoluene (BHT) solution, and 500 µL of 30% trichloroacetic acid (TCA). The mixture was vortexed and kept on ice for 2 hours before being centrifuged at 2000 rpm for 15 minutes. From the supernatant, 1 mL was transferred to another tube, and 75 µL of EDTA and 250 µL of TBA were added. The tubes were vortexed and incubated in a hot water bath for 15 minutes. After cooling to room temperature, the absorbance was measured at 532 nm using a UV-Vis spectrophotometer. The MDA concentration was calculated using the formula:

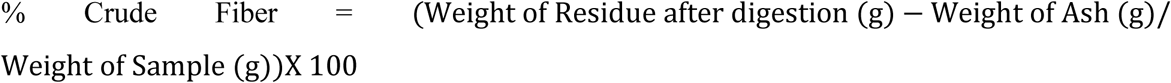

Where F represents the dilution factor, 6.41 is the extinction coefficient, and A is the absorbance.

## STATISTICAL ANALYSIS

All data were expressed as the mean of six replicates ± standard error of the mean (S.E.M) (except for the proximate and amino acid content analysis where three replicates were used). Statistical evaluation of data was performed using SPSS version 27.0 using one-way analysis of variance (ANOVA). Followed by Duncan’s posthoc test for multiple comparisons. Values were considered statistically significant at p ≤ 0.05 (confidence level = 95%).

## RESULTS

**Table 2:** Proximate analyses result (%) of the experimental feeds used. Results are expressed as Mean ±S.D, (n=3). ANOVA analysis was carried out using Duncan’s post hoc test for multiple comparisons. Superscripts with different letters show statistical differences. Values were considered statistically significant at p ≤ 0.05 (confidence level = 95%).

| Sample | Crude Protein (%) | Moisture (%) | Fat (%) | Ash (%) | Carbohydrate (%) |
| --- | --- | --- | --- | --- | --- |
| Regular Animal Chow | 10.19<sup>b</sup>±0.28 | 6.44<sup>b</sup>±0.23 | 4.21<sup>b</sup>±0.11 | 1.83<sup>b</sup>±0.03 | 74.61<sup>c</sup>±0.65 |
| Milled Maize | 9.68<sup>b</sup>±0.21 | 7.81<sup>b</sup>±0.13 | 4.39<sup>b</sup>±0.13 | 1.68<sup>b</sup>±0.04 | 79.15<sup>c</sup>±0.58 |
| Red Sorghum | 5.39<sup>a</sup>±0.25 | 5.31<sup>a</sup>±0.18 | 3.31<sup>a</sup>±0.07 | 1.62<sup>a</sup>±0.02 | 84.37<sup>c</sup>±0.61 |
| White Sorghum | 5.54<sup>a</sup>±0.19 | 5.57<sup>a</sup>±0.17 | 3.9<sup>b</sup>±0.14 | 1.61<sup>a</sup>±0.03 | 83.38<sup>c</sup>±0.51 |

**Table 3:**
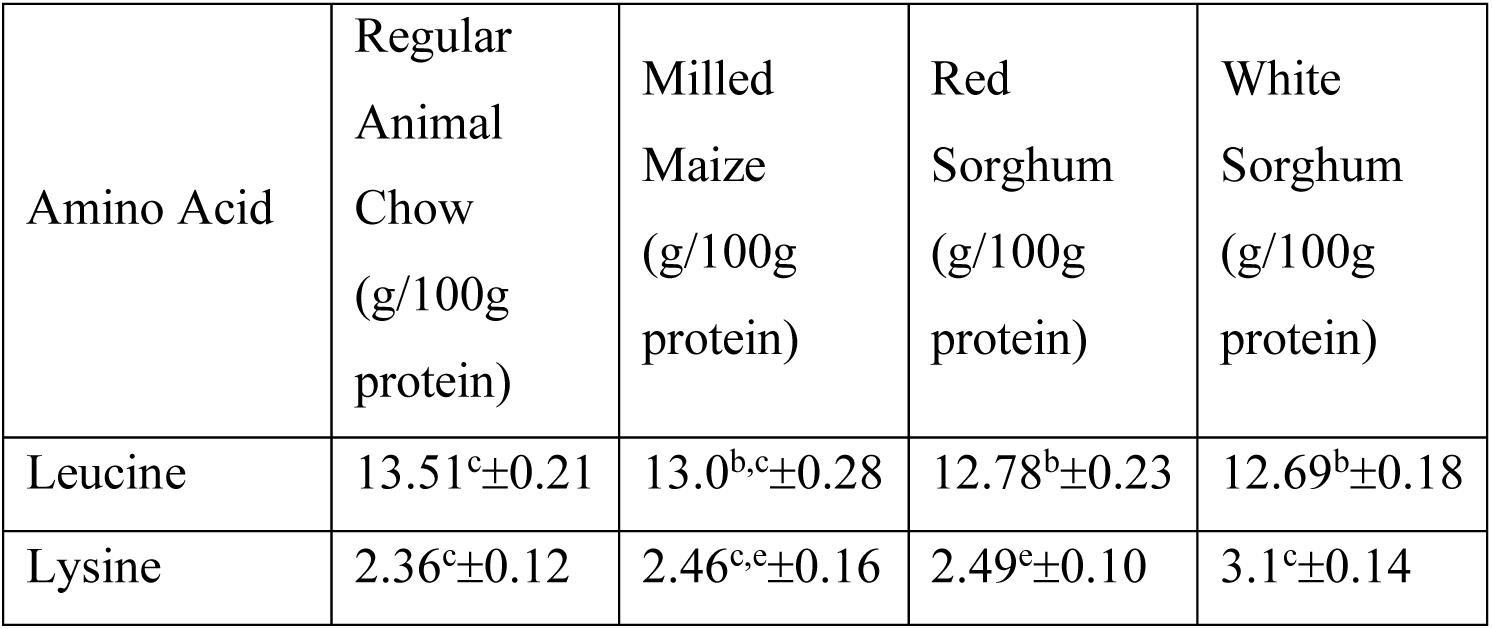
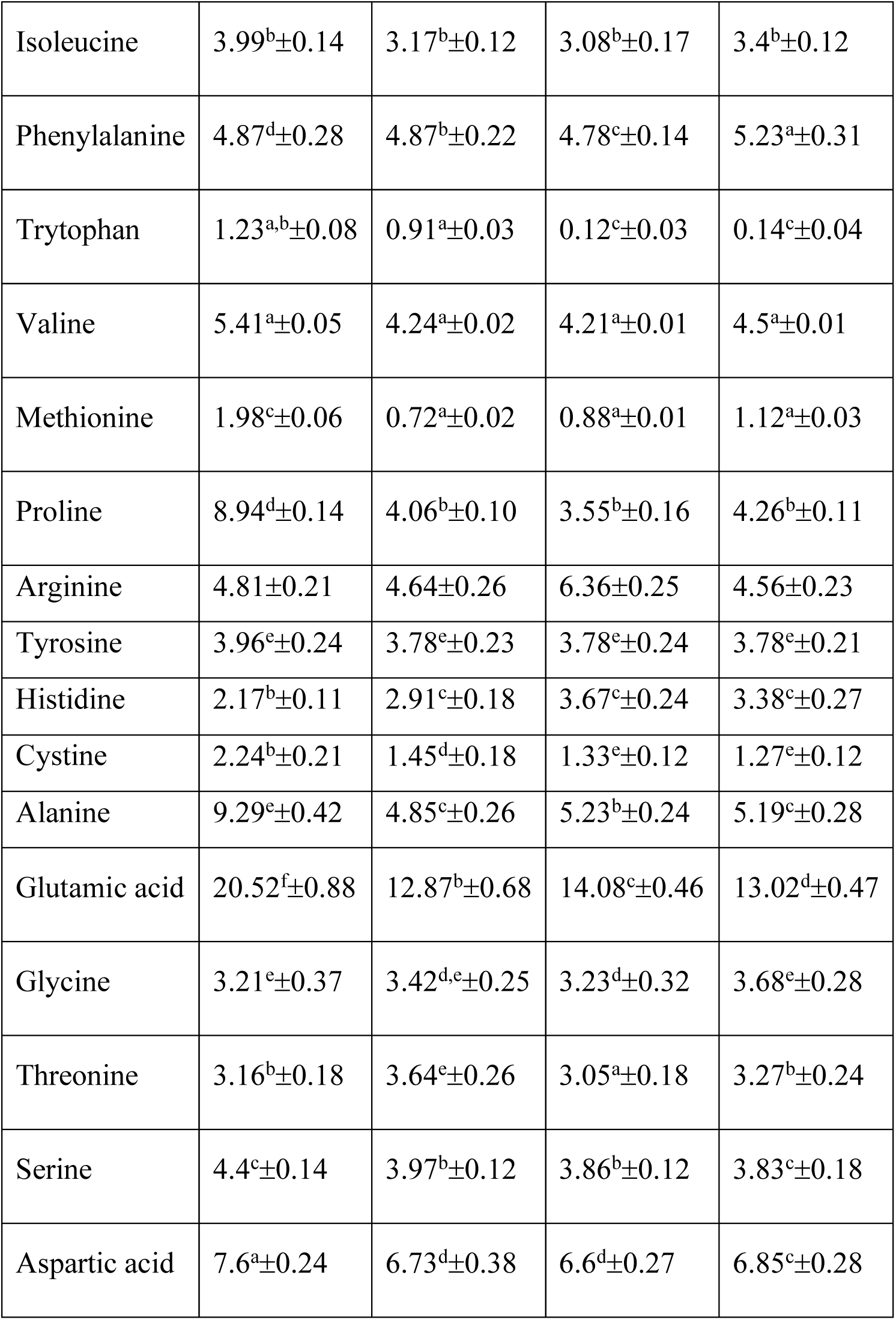
Total Individual Amino Acids Content in Experimental Feeds. Results are expressed as Mean ±S.D, (n=3). ANOVA analysis was carried out using Duncan’s post hoc test for multiple comparisons. Superscripts with different letters show statistical differences. Values were considered statistically significant at p ≤ 0.05 (confidence level = 95%).

**Table 4:** Serum testosterone concentration (ng/dL) over time in the experimental groups. Results are expressed as mean ±S.E.M, (n=6). ANOVA analysis was carried out using Duncan’s post hoc test for multiple comparisons. Superscripts with different letters show statistical differences. Values were considered statistically significant at p ≤ 0.05 (confidence level = 95%).

| Group | Testosterone<br>level Day 1<br>(ng/dL) | Testosterone<br>level Day 4<br>(ng/dL) | Testosterone<br>level Day 7<br>(ng/dL) | Testosterone<br>level Day 10<br>(ng/dL) | Testosterone<br>level Day 13<br>(ng/dL) |
| --- | --- | --- | --- | --- | --- |
| A | 0.648<sup>c</sup>±0.025 | 0.509<sup>b</sup>±0.021 | 0.612<sup>b</sup>±0.037 | 0.536<sup>a</sup>±0.052 | 0.651<sup>a</sup>±0.028 |
| B | 0.625<sup>c</sup>±0.031 | 0.622<sup>b</sup>±0.047 | 0.599<sup>b</sup>±0.084 | 0.581<sup>a</sup>±0.027 | 0.692<sup>b</sup>±0.083 |
| C | 0.856<sup>b</sup>±0.029 | 0.822<sup>c</sup>±0.037 | 0.834<sup>c</sup>±0.022 | 0.818<sup>c</sup>±0.092 | 0.824<sup>c</sup>±0.038 |
| D | 0.884<sup>b</sup>±0.048 | 0.832<sup>c</sup>±0.042 | 1.325<sup>a</sup>±0.087 | 1.619<sup>b</sup>±0.095 | 1.556<sup>b</sup>±0.097 |
| E | 0.977<sup>a</sup>±0.029 | 1.112<sup>d</sup>±0.078 | 1.254<sup>c</sup>±0.193 | 1.992<sup>d</sup>±0.129 | 1.678<sup>d</sup>±0.074 |
| F | 0.509<sup>c</sup>±0.083 | 0.556<sup>a</sup>±0.028 | 0.486<sup>e</sup>±0.098 | 0.382<sup>e</sup>±0.049 | 0.324<sup>e</sup>±0.028 |
| G | 0.667<sup>c</sup>±0.027 | 0.627<sup>b</sup>±0.022 | 0.632<sup>d</sup>±0.037 | 0.564<sup>a</sup>±0.038 | 0.483<sup>g</sup>±0.033 |
| H | 0.934<sup>b</sup>±0.038 | 0.884<sup>e</sup>±0.029 | 0.888<sup>c</sup>±0.084 | 0.832<sup>i</sup>±0.035 | 0.796<sup>f</sup>±0.084 |
| I | 0.882<sup>b</sup>±0.094 | 0.434<sup>c</sup>±0.085 | 0.328<sup>a</sup>±0.028 | 0.253<sup>e</sup>±0.019 | 0.283<sup>e</sup>±0.023 |
| J | 0.912<sup>a</sup>±0.034 | 0.523<sup>b</sup>±0.038 | 0.487<sup>e</sup>±0.081 | 0.345<sup>d</sup>±0.094 | 0.319<sup>d</sup>±0.038 |

**Table 5:**
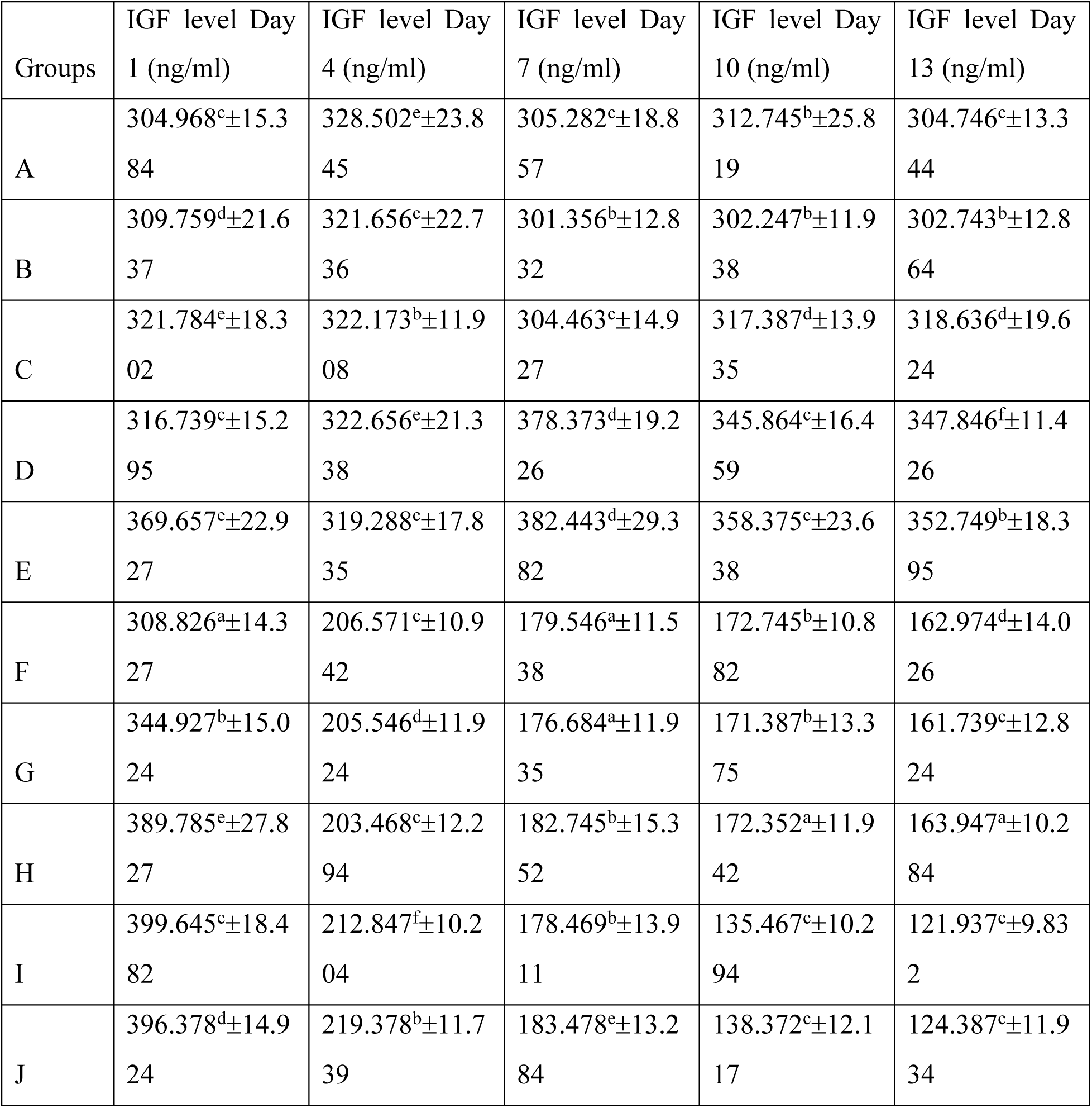
Serum Insulin-like Growth Factor (IGF) concentration (ng/ml) over time in the experimental groups. Results are expressed as mean ±S.E.M, (n=6). ANOVA analysis was carried out using Duncan’s post hoc test for multiple comparisons. Superscripts with different letters show statistical differences. Values were considered statistically significant at p ≤ 0.05 (confidence level = 95%).

**Table 6:**
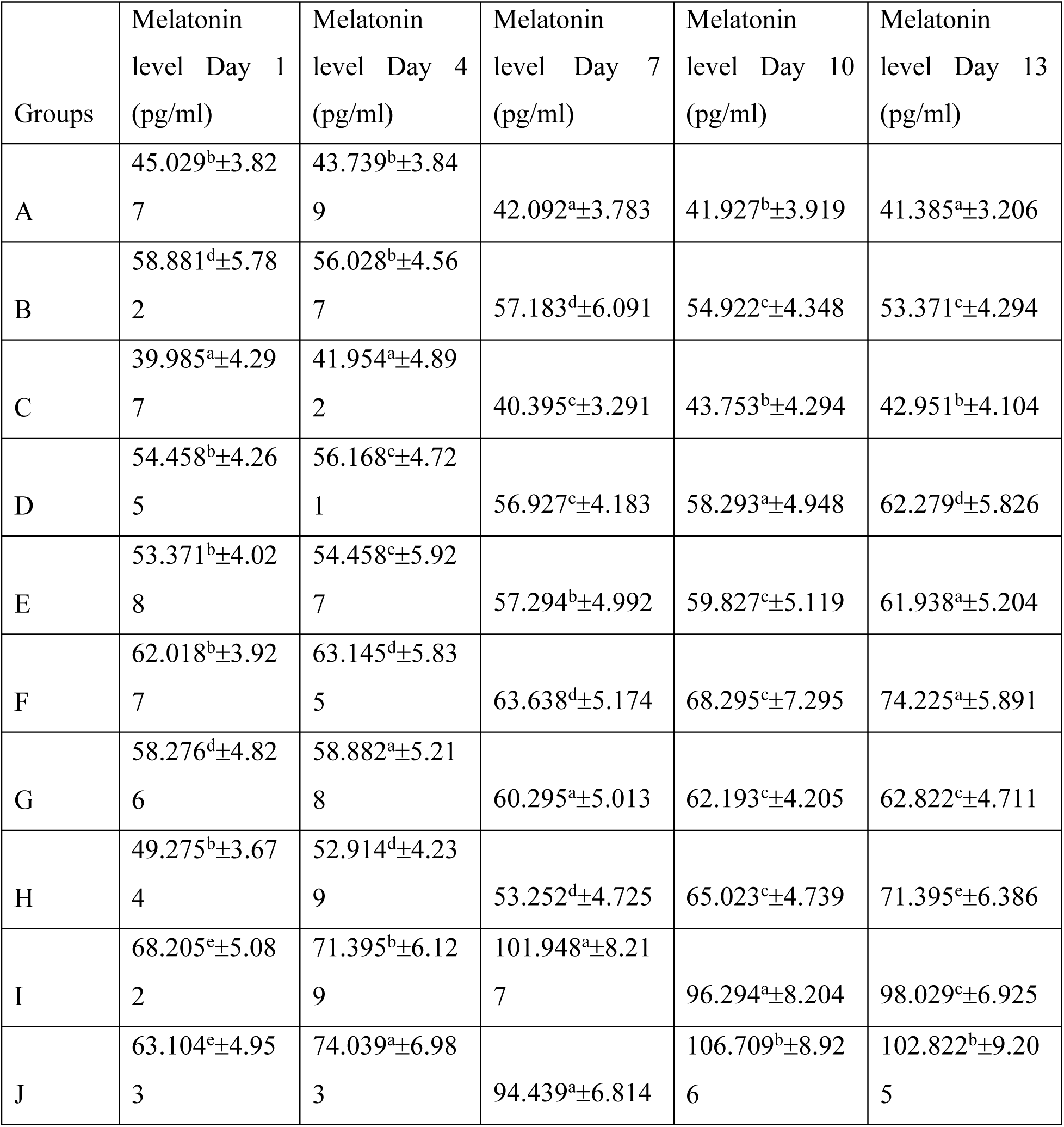
Serum melatonin concentration (pg/ml) over time in the experimental groups. Results are expressed as mean ±S.E.M, (n=6). ANOVA analysis was carried out using Duncan’s post hoc test for multiple comparisons. Superscripts with different letters show statistical differences. Values were considered statistically significant at p ≤ 0.05 (confidence level = 95%).

**Table 7:**
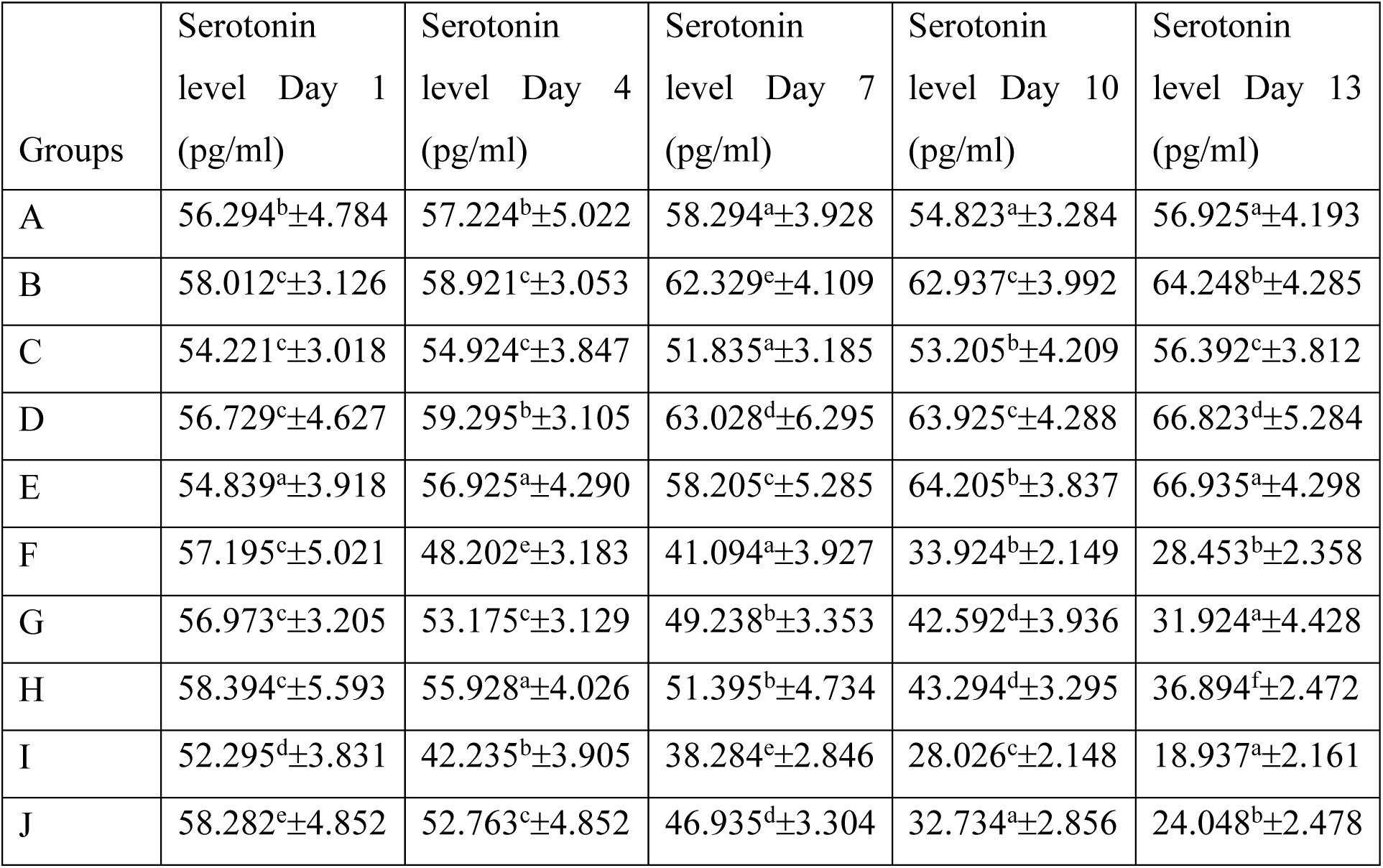
Serum serotonin concentration (pg/ml) over time in the experimental groups. Results are expressed as mean ±S.E.M, (n=6). ANOVA analysis was carried out using Duncan’s post hoc test for multiple comparisons. Superscripts with different letters show statistical differences. Values were considered statistically significant at p ≤ 0.05 (confidence level = 95%).

**Table 8:**
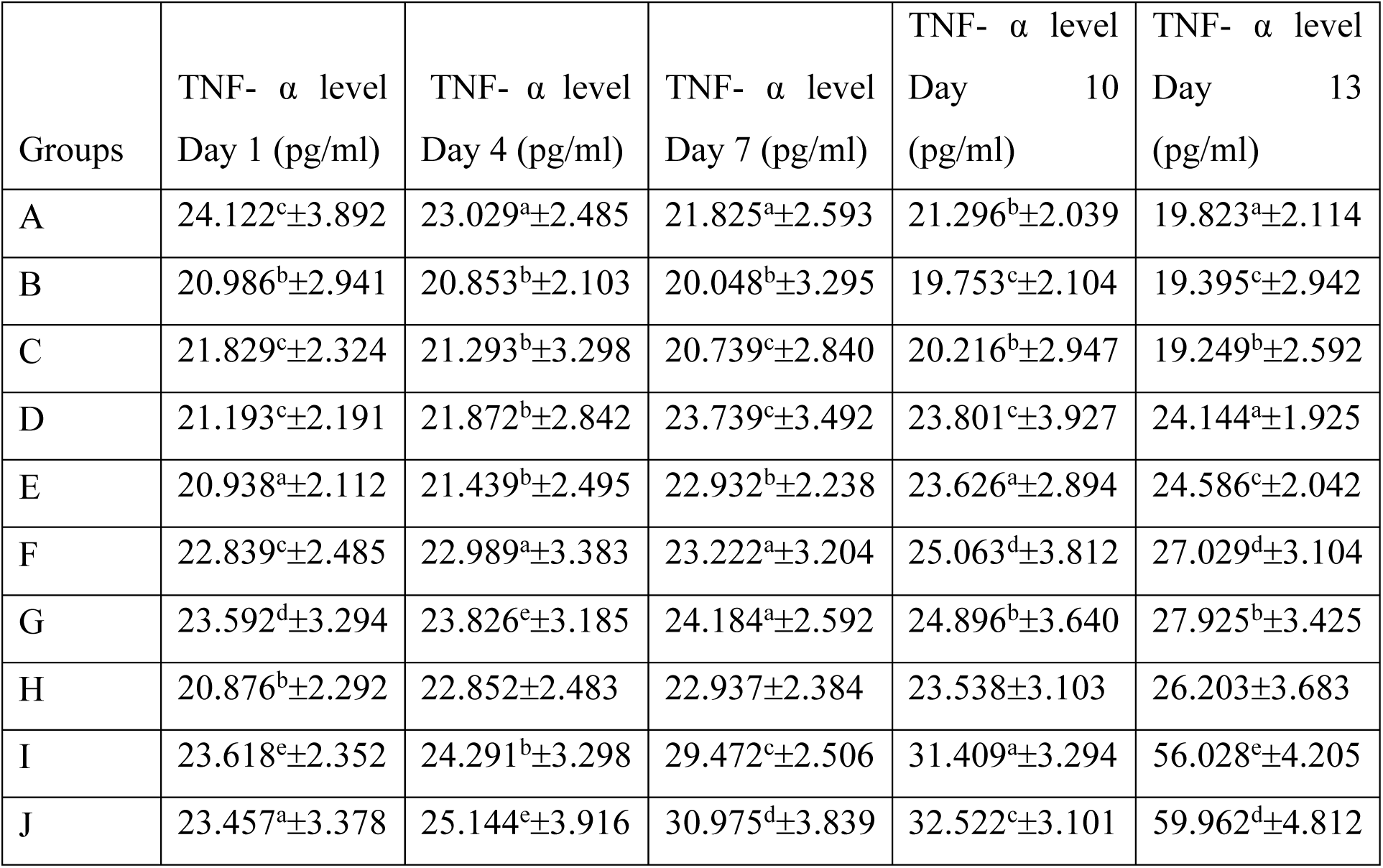
Serum Tumor Necrosis Factor-alpha (TNF-α) concentration (pg/ml) over time in the experimental groups. Results are expressed as mean ±S.E.M, (n=6). ANOVA analysis was carried out using Duncan’s post hoc test for multiple comparisons. Superscripts with different letters show statistical differences. Values were considered statistically significant at p ≤ 0.05 (confidence level = 95%).

**Table 9:**
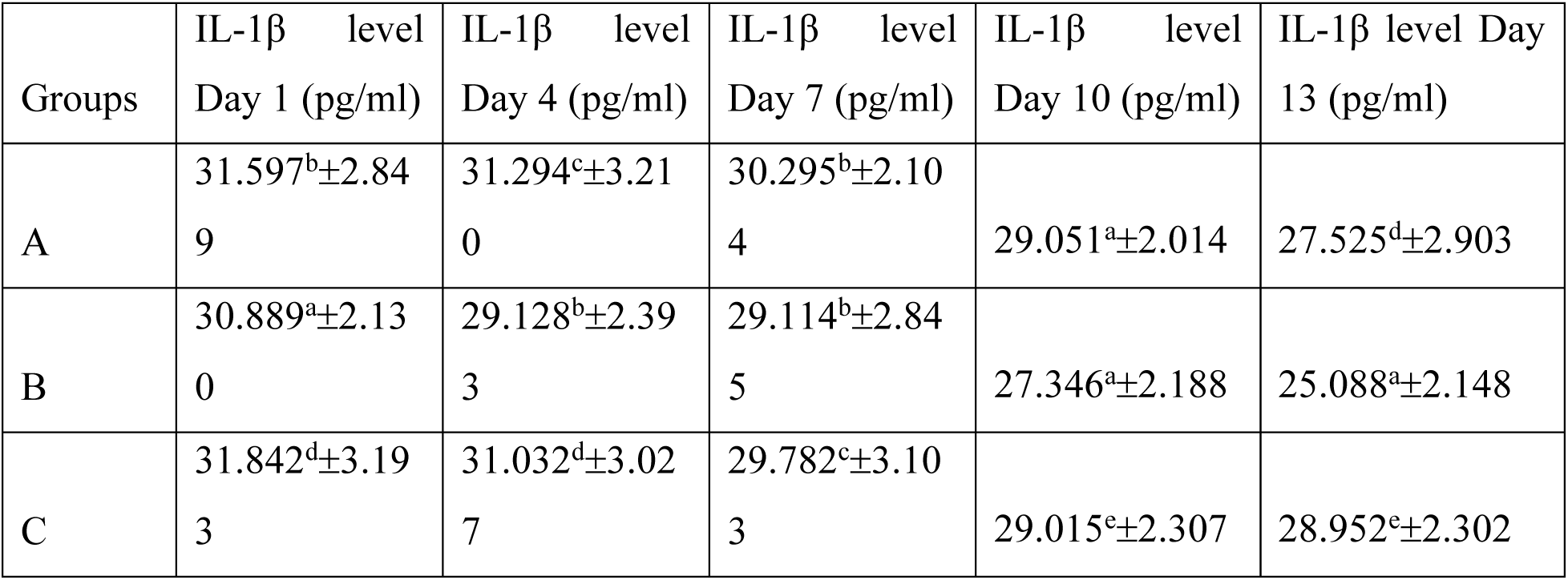
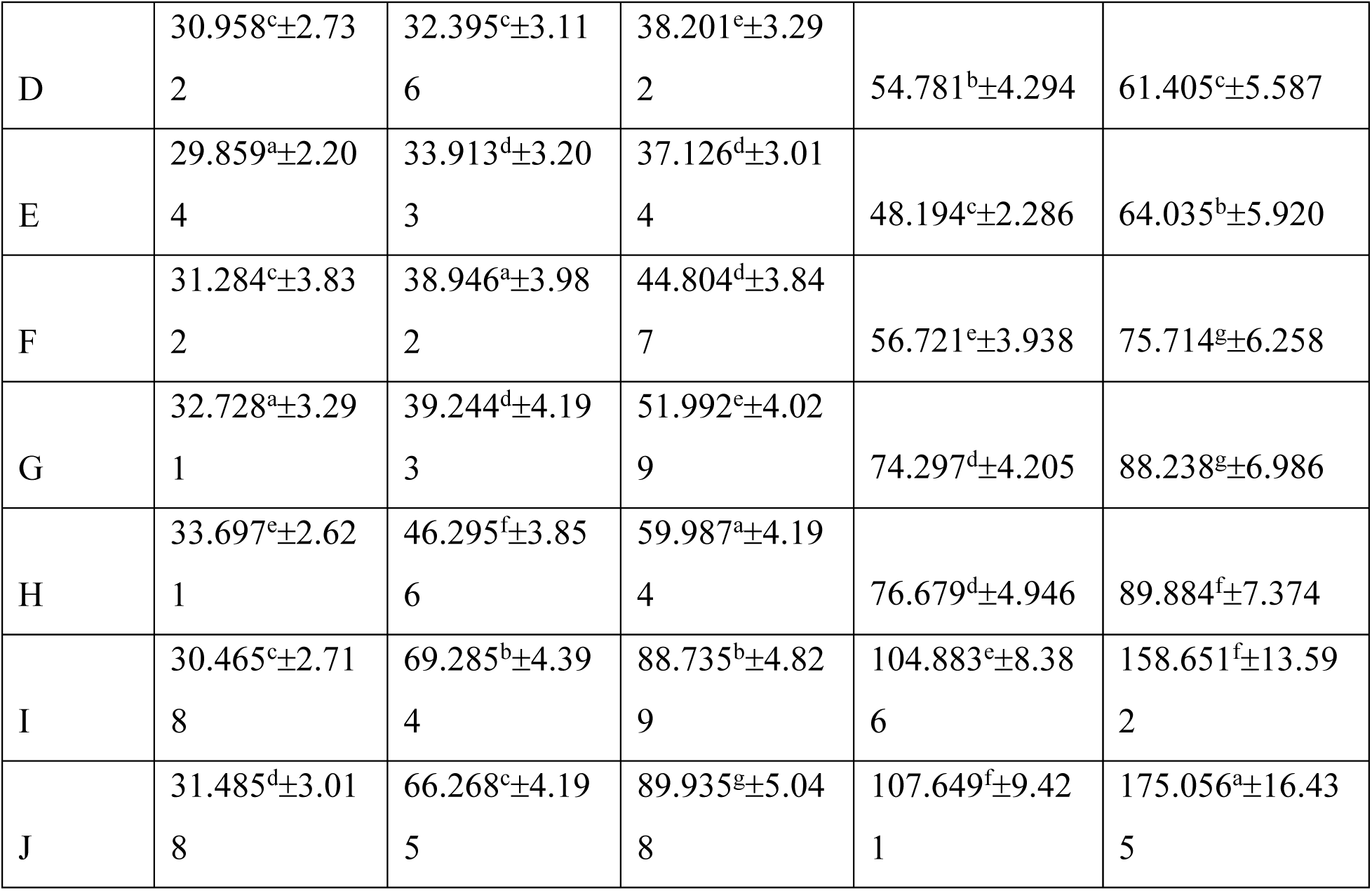
Serum Interleukin-1 beta (IL-1β) concentration (pg/ml) over time in the experimental groups. Results are expressed as mean ±S.E.M, (n=6). ANOVA analysis was carried out using Duncan’s post hoc test for multiple comparisons. Superscripts with different letters show statistical differences. Values were considered statistically significant at p ≤ 0.05 (confidence level = 95%).

**Table 10:** Serum Superoxide Dismutase (SOD) activity (U/L) over time in the experimental groups. Results are expressed as mean ±S.E.M, (n=6). ANOVA analysis was carried out using Duncan’s post hoc test for multiple comparisons. Superscripts with different letters show statistical differences. Values were considered statistically significant at p ≤ 0.05 (confidence level = 95%).

| Groups | SOD (U/L)<br>Day 1 | SOD (U/L)<br>Day 4 | SOD (U/L)<br>Day 7 | SOD (U/L)<br>Day 10 | SOD (U/L)<br>Day 13 |
| --- | --- | --- | --- | --- | --- |
| A | 1.192<sup>e</sup>±0.028 | 1.349<sup>e</sup>±0.031 | 1.428<sup>b</sup>±0.028 | 1.849<sup>c</sup>±0.016 | 2.257<sup>c</sup>±0.069 |
| B | 1.139<sup>b</sup>±0.013 | 1.364<sup>d</sup>±0.023 | 1.578<sup>c</sup>±0.039 | 1.528<sup>c</sup>±0.023 | 1.984<sup>d</sup>±0.035 |
| C | 1.060<sup>e</sup>±0.008 | 1.118<sup>b</sup>±0.025 | 1.342<sup>c</sup>±0.018 | 1.697<sup>d</sup>±0.027 | 1.858<sup>d</sup>±0.052 |
| D | 1.243<sup>a</sup>±0.021 | 1.01<sup>d</sup>±0.016 | 0.816<sup>c</sup>±0.005 | 0.729<sup>b</sup>±0.012 | 0.732<sup>c</sup>±0.006 |
| E | 1.155<sup>b</sup>±0.019 | 0.981<sup>e</sup>±0.007 | 0.732<sup>d</sup>±0.007 | 0.619<sup>f</sup>±0.008 | 0.666<sup>f</sup>±0.004 |
| F | 1.281<sup>b</sup>±0.024 | 0.975<sup>d</sup>±0.003 | 0.894<sup>e</sup>±0.004 | 0.788<sup>c</sup>±0.006 | 0.872<sup>c</sup>±0.008 |
| G | 1.088<sup>d</sup>±0.009 | 1.021<sup>e</sup>±0.009 | 0.928<sup>a</sup>±0.007 | 0.854<sup>b</sup>±0.014 | 0.731<sup>c</sup>±0.009 |
| H | 1.142<sup>a</sup>±0.011 | 1.042<sup>b</sup>±0.006 | 0.946<sup>b</sup>±0.008 | 0.982<sup>e</sup>±0.008 | 0.799<sup>d</sup>±0.005 |
| I | 1.238<sup>c</sup>±0.029 | 1.038<sup>a</sup>±0.013 | 0.911<sup>b</sup>±0.010 | 0.513<sup>e</sup>±0.006 | 0.127<sup>d</sup>±0.003 |
| J | 1.034<sup>e</sup>±0.014 | 0.994<sup>d</sup>±0.008 | 0.863<sup>a</sup>±0.008 | 0.489<sup>c</sup>±0.004 | 0.212<sup>b</sup>±0.004 |

**Table 11:** Serum Catalase (CAT) activity (U/L) over time in the experimental groups. Results are expressed as mean ±S.E.M, (n=6). ANOVA analysis was carried out using Duncan’s post hoc test for multiple comparisons. Superscripts with different letters show statistical differences. Values were considered statistically significant at p ≤ 0.05 (confidence level = 95%).

| Groups | CAT (U/L)<br>Day 1 | CAT (U/L)<br>Day 4 | CAT (U/L)<br>Day 7 | CAT (U/L)<br>Day 10 | CAT (U/L)<br>Day 13 |
| --- | --- | --- | --- | --- | --- |
| A | 16.396<sup>b</sup>±1.053 | 16.294<sup>c</sup>±1.948 | 16.253<sup>c</sup>±1.836 | 15.867<sup>b</sup>±1.304 | 15.938<sup>b</sup>±1.305 |
| B | 15.358<sup>a</sup>±1.397 | 15.283<sup>a</sup>±1.437 | 15.038<sup>c</sup>±1.204 | 15.184<sup>b</sup>±1.743 | 15.395<sup>b</sup>±1.593 |
| C | 13.034<sup>d</sup>±1.830 | 13.402<sup>e</sup>±1.039 | 13.257<sup>b</sup>±1.958 | 13.028<sup>c</sup>±1.483 | 13.635<sup>b</sup>±1.942 |
| D | 15.532<sup>d</sup>±1.932 | 15.082<sup>d</sup>±1.842 | 13.997<sup>b</sup>±1.437 | 13.284<sup>b</sup>±1.290 | 13.018<sup>b</sup>±1.224 |
| E | 14.924<sup>a</sup>±1.527 | 14.284<sup>b</sup>±1.338 | 14.028<sup>c</sup>±1.753 | 13.392<sup>b</sup>±1.394 | 13.149<sup>b</sup>±1.685 |
| F | 13.883<sup>b</sup>±1.068 | 13.193<sup>c</sup>±1.749 | 12.485<sup>e</sup>±1.458 | 12.294<sup>d</sup>±1.479 | 11.692<sup>a</sup>±1.453 |
| G | 14.924<sup>c</sup>±1.944 | 13.989<sup>a</sup>±1.832 | 13.341<sup>b</sup>±1.579 | 12.038<sup>f</sup>±1.120 | 11.839<sup>e</sup>±1.892 |
| H | 13.098<sup>d</sup>±1.822 | 12.395<sup>f</sup>±1.136 | 10.395<sup>c</sup>±1.021 | 9.843<sup>a</sup>±1.092 | 7.659<sup>b</sup>±0.429 |
| I | 17.165<sup>b</sup>±1.643 | 12.114<sup>e</sup>±1.285 | 10.145<sup>a</sup>±1.038 | 9.246<sup>c</sup>±1.142 | 8.386<sup>c</sup>±0.822 |
| J | 16.827<sup>a</sup>±1.750 | 13.283<sup>e</sup>±1.489 | 11.294<sup>c</sup>±1.028 | 8.518<sup>b</sup>±1.038 | 7.576<sup>d</sup>±0.338 |

**Table 12:**
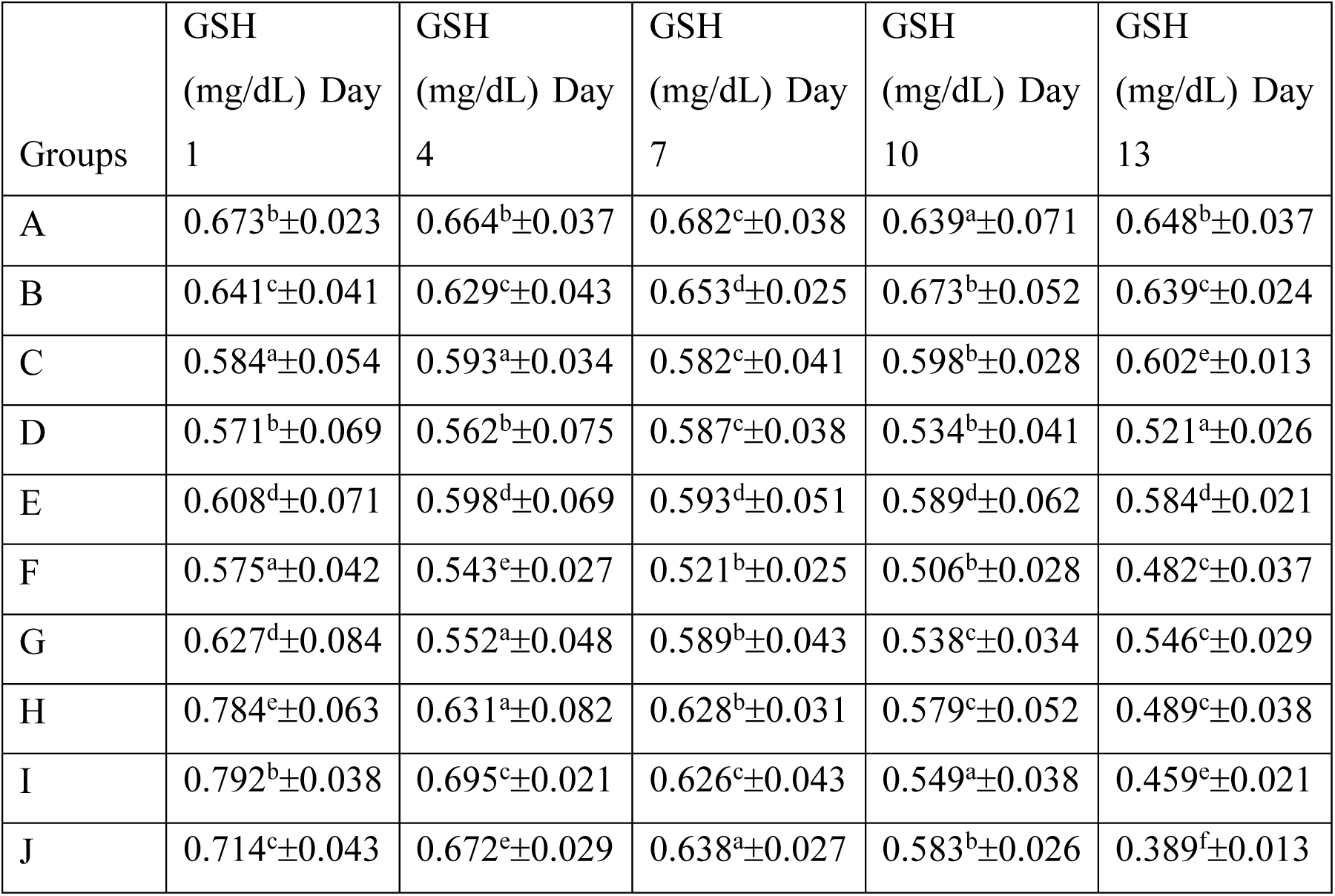
Glutathione (GSH) concentration (mg/dL) over time in the experimental groups. Results are expressed as mean±S.E.M, (n=6). ANOVA analysis was carried out using Duncan’s post hoc test for multiple comparisons. Superscripts with different letters show statistical differences. Values were considered statistically significant at p ≤ 0.05 (confidence level = 95%).

**Table 13:** Malondialdehyde (MDA) concentration (µmol/L) over time in the experimental groups. Results are expressed as mean±S.E.M, (n=6). ANOVA analysis was carried out using Duncan’s post hoc test for multiple comparisons. Superscripts with different letters show statistical differences. Values were considered statistically significant at p ≤ 0.05 (confidence level = 95%).

| Groups | MDA<br>($\mu\text{mol/L}$)<br>Day 1 | MDA<br>($\mu\text{mol/L}$)<br>Day 4 | MDA<br>($\mu\text{mol/L}$)<br>Day 7 | MDA<br>($\mu\text{mol/L}$)<br>Day 10 | MDA<br>($\mu\text{mol/L}$) Day<br>13 |
| --- | --- | --- | --- | --- | --- |
| A | 3.944<sup>b</sup>±0.285 | 4.082<sup>b</sup>±0.383 | 4.285<sup>a</sup>±0.392 | 4.184<sup>c</sup>±0.284 | 4.358<sup>b</sup>±0.293 |
| B | 4.964<sup>d</sup>±0.434 | 5.294<sup>a</sup>±0.492 | 5.026<sup>b</sup>±0.441 | 5.287<sup>c</sup>±0.623 | 5.321<sup>c</sup>±0.550 |
| C | 4.352<sup>a</sup>±0.236 | 4.736<sup>c</sup>±0.424 | 4.392<sup>a</sup>±0.313 | 4.928<sup>c</sup>±0.431 | 5.015<sup>c</sup>±0.442 |
| D | 4.709<sup>b</sup>±0.411 | 4.923<sup>c</sup>±0.393 | 5.184<sup>a</sup>±0.492 | 5.867<sup>d</sup>±0.592 | 6.801<sup>c</sup>±0.394 |
| E | 4.403<sup>b</sup>±0.375 | 4.794<sup>a</sup>±0.284 | 5.069<sup>a</sup>±0.401 | 5.837<sup>d</sup>±0.423 | 6.239<sup>c</sup>±0.483 |
| F | 3.896<sup>d</sup>±0.193 | 4.029<sup>c</sup>±0.385 | 4.848<sup>a</sup>±0.331 | 6.487<sup>e</sup>±0.595 | 8.928<sup>e</sup>±0.624 |
| G | 4.301<sup>a</sup>±0.352 | 4.846<sup>c</sup>±0.385 | 5.729<sup>f</sup>±0.378 | 7.027<sup>d</sup>±0.628 | 8.184<sup>b</sup>±0.388 |
| H | 4.148<sup>e</sup>±0.420 | 5.823<sup>b</sup>±0.332 | 7.218<sup>d</sup>±0.417 | 8.639<sup>d</sup>±0.491 | 9.029<sup>c</sup>±0.596 |
| I | 4.352<sup>a</sup>±0.391 | 5.938<sup>d</sup>±0.284 | 6.839<sup>e</sup>±0.433 | 8.478<sup>g</sup>±0.783 | 9.882<sup>f</sup>±0.652 |
| J | 4.037<sup>b</sup>±0.283 | 5.892<sup>f</sup>±0.384 | 7.039<sup>e</sup>±0.482 | 8.932<sup>c</sup>±0.682 | 10.984<sup>d</sup>±0.835 |

**Figure 1:**
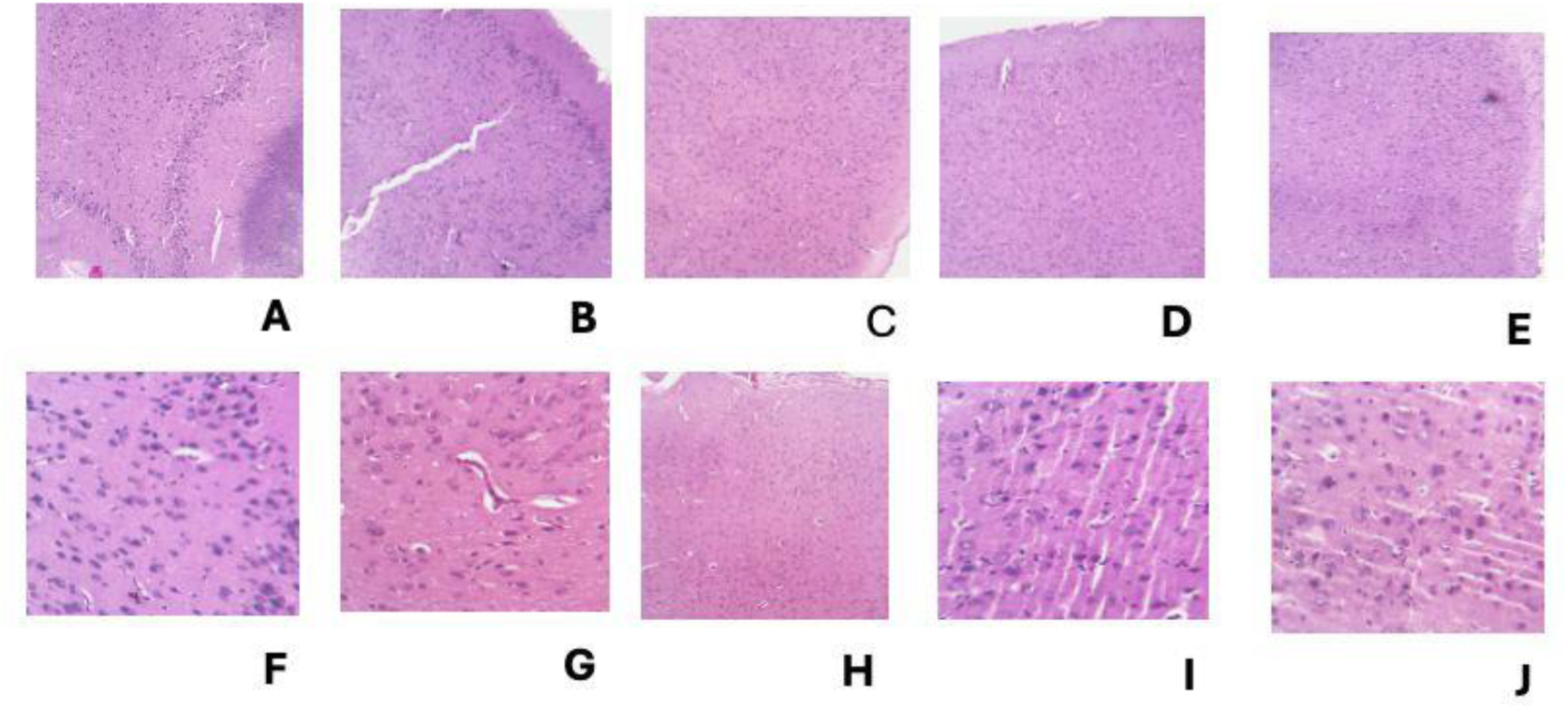
Section shows multilayers of cerebrum in the experimental groups composed of the outer molecular layer, followed by the outer granular layer, outer plexiform layer, inner granular cell layer, inner plexiform layer, and pleomorphic layer in sequential order. The cells are seen in the background of brain parenchyma The first hypocellular area is known as the molecular layer. The outer granular layers are composed of a linear layer of densely populated cells beneath the hypocellular molecular cell. The pleomorphic cells are large cells with prominent nucleoli and abundant cytoplasm in groups F-J.

## DISCUSSION

This study explores how sleep deprivation, caffeine intake, and diet affect biochemical biomarkers involved in growth, sleep, inflammation, and oxidative stress. Amino acid composition and proximate analysis of the experimental diets revealed that maize had the highest tryptophan and protein comparable to the standard rat chow, although significant differences exist. Sorghum, both white and red, have lower tryptophan composition and protein content, although they have a higher lysine, histidine (g/100g protein), and carbohydrates (%). This corroborates the findings by (Pan et al., 2021) who reported a lower tryptophan and protein composition in a sorghum-based diet compared to a corn-based diet, potentially influencing sleep-related biomarkers negatively.

The analysis of serum testosterone levels across various experimental groups reveals significant insights into the interactions between diet, caffeine intake, sleep deprivation, and hormonal balance. Groups A, B, and C, which were housed in regular cages without caffeine, exhibited relatively stable testosterone levels. This stability suggests minimal impact under non-stressful conditions, aligning with previous research indicating that stable housing and a balanced diet are crucial for maintaining hormonal homeostasis (Smith et al., 2019). Notably, Group E, which was subjected to a sorghum diet and caffeine, experienced a significant rise in testosterone levels, peaking on Day 10. This increase could be attributed to the combined effects of tryptophan-rich sorghum and the stimulatory influence of caffeine. This finding is consistent with studies by (Donald et al., 2017) who reported that caffeine can enhance testosterone levels when accompanied by adequate dietary support, although the long-term effects were not assessed.

In contrast, Groups F to J, which were housed in disk-over-water cages indicative of sleep deprivation, demonstrated sharp declines in both testosterone and insulin-like growth factor (IGF) levels. Particularly in Groups I and J, which also received caffeine, the marked hormonal decrease suggests a compounded negative effect of sleep deprivation and caffeine, supporting findings by (Van Cauter et al., 2008) that highlighted similar disruptions in hormonal balance under sleep-deprived conditions. Group D, which had a maize diet and caffeine but no sleep deprivation, initially exhibited a rise in testosterone levels that later escalated significantly, possibly due to caffeine’s role in increasing testosterone in non-sleep-deprived animals, as noted by (Park et al., 2015). Correspondingly, IGF levels followed a similar trajectory, showing significant drops in sleep-deprived groups, particularly those with caffeine, indicating a stress-induced catabolic state. This observation corroborates findings from (Spiegel et al., 2005), who underscored the negative impact of chronic stress on IGF levels.

Furthermore, the serum testosterone analysis indicated that caffeine intake and a tryptophan-deficient diet significantly impact testosterone levels, suggesting a reduced sex drive. This reduction may contribute to observed behaviors such as loss of fur, irritability, and loss of muscle function in experimental rats, as testosterone levels decreased throughout the duration of the experiment. A pronounced reduction in testosterone was particularly evident in rats housed in disk-over-water cages, with the most significant declines observed in Groups I and J, which received a combination of daily caffeine administration (20 mg/kg body weight) and were housed in these cages. Statistical significance in the reduction was noted for Groups G and H, which were also housed in disk-over-water cages during the experimental period, compared to those in normal cages and those in normal cages receiving 20 mg/kg body weight caffeine (Groups D and E). Conversely, Groups D and E demonstrated a marked increase in testosterone levels when caffeine was administered in normal cages over 13 days. The observed decline in testosterone levels among caffeine-administered and tryptophan-deficient groups aligns with findings by (Eliot, 2021), who reported that caffeine and sleep deprivation can impair testosterone synthesis, resulting in behavioral and physiological changes such as aggression, muscle atrophy, and hair loss.

The analysis of serum IGF-1 levels further highlights the detrimental effects of sleep deprivation and caffeine administration. A drastic decrease in IGF-1 levels was observed in groups subjected to sleep deprivation using the disk-over-water cage or a combination of both caffeine administration and disk-over-water housing, indicating potential growth hormone deficiency and malnutrition. This decrease in serum IGF-1 in sleep-deprived groups echoes research by (Wan et al., 2022) which found that sleep deprivation impairs growth factor production, compounded by caffeine’s antagonistic effects on growth hormone secretion. Interestingly, the trend observed in rats from Groups D and E showed an increase in serum IGF-1 concentrations as the duration of caffeine administration lengthened, suggesting a change in physiological condition and increased growth hormone secretion. This finding challenges the previously reported antagonistic effect of caffeine on growth hormone secretion noted by (Wan et al., 2022).

Analysis of serum melatonin levels revealed a steady decrease in the control groups A and B across the experimental days, except for Group C, which exhibited a gradual increase. This rise in Group C may be attributed to the intake of tryptophan-deficient sorghum. In contrast, a marked increase in serum melatonin levels was observed in all sleep-deprived groups (D-J), particularly in Groups I and J, where rats experienced both caffeine administration (20 mg/kg) and housing in disk-over-water cages. The significant elevation in melatonin levels suggests that the experimental rats may have experienced disrupted circadian rhythms, evidenced by their lack of vigor and pronounced behavioral changes, such as massive fur loss and delirium in Groups I and J.

Serum serotonin analysis showed stable levels across the control groups, with a peak observed in Groups D and E, which received caffeine while housed in regular cages. However, a decrease in serotonin levels was evident in all groups (F-J) subjected to sleep deprivation, particularly in Groups I and J. These fluctuations in melatonin and serotonin suggest that the combination of sleep deprivation and caffeine intake disrupts normal circadian rhythms, as similarly noted by (Jenkins et al., 2016), where melatonin suppression and serotonin depletion were linked to mood disorders and cognitive deficits.

The analysis of serum pro-inflammatory markers TNF-α and IL-1β showed a steady decrease in control groups but an increase in all sleep-deprived groups, with the highest levels observed in rats receiving daily caffeine administration and housed in disk-over-water cages. This increase in inflammatory markers in Groups I and J underscores the systemic inflammation induced by chronic sleep loss, which is exacerbated by caffeine. Correspondingly, the serum superoxide dismutase (SOD) activity revealed a steady increase in the control groups, while a sharp reduction was noted in sleep-deprived groups, particularly in Groups I and J. Additionally, catalase, glutathione peroxidase activity, and glutathione levels demonstrated a statistically insignificant decrease in the control groups (A-E) but showed significant reductions in those housed in disk-over-water cages (F-J) alongside caffeine administration. These findings are consistent with previous studies indicating that chronic sleep deprivation can lead to systemic inflammation and increased oxidative stress, as reported by (Irwin et al., 2016).

An increase in malondialdehyde (MDA) levels across all experimental groups was observed, with the highest increases in those housed in disk-over-water cages. This trend may indicate oxidative stress, tissue damage, and chronic inflammatory conditions, particularly given the erratic behavior, social anxiety, and muscle function loss observed in the experimental rats. The combination of these stressors, evidenced by elevated MDA levels, suggests significant oxidative damage that could explain the severe behavioral and physiological deterioration, aligning with models of neuroinflammation and oxidative stress described by (Bellesi et al., 2017).

Histomorphological examination revealed no abnormalities in light microscopy of the fixed brain tissues in the cerebrum for most experimental groups. However, Groups I and J exhibited pleomorphic cells characterized by large sizes, prominent nucleoli, and abundant cytoplasm, indicating inflammation, malignancies, and dysplasia. These histopathological changes, particularly in groups exposed to both sleep deprivation and high caffeine intake, point to potential neurodegenerative effects, consistent with existing literature on the compounded impact of these stressors on brain health.

Interestingly, lower doses of caffeine have been shown to exhibit neuroprotective effects, enhancing cognitive function and reducing oxidative stress markers. In contrast, higher doses, like those used in this study (20 mg/kg), are more likely to exacerbate stress responses and contribute to neuroinflammation (Sousa et al., 2018). This highlights the importance of dosage when considering caffeine’s effects on physiological and psychological health.

Moreover, studies focusing on the neuroprotective effects of antioxidant-rich diets, such as the Mediterranean diet or those high in omega-3 fatty acids, suggest that these dietary patterns can significantly buffer the negative effects of stress and high caffeine intake. For example, dietary supplementation with omega-3 fatty acids has been shown to reduce levels of inflammatory cytokines (such as TNF-α and IL-1β) and oxidative stress markers that were elevated in this study under conditions of high caffeine and sleep deprivation. Additionally, diets rich in polyphenols, found in fruits and vegetables, have been shown to enhance brain-derived neurotrophic factor (BDNF) expression, providing neuroprotection even under chronic stress conditions (Fernando, 2008).

### Conclusion

This study highlights the pivotal role of sleep quality in maintaining hormonal balance, demonstrating that while caffeine can enhance testosterone levels under non-stressful conditions, its effects become detrimental in the context of sleep deprivation. The findings offer significant insights into the interactions between dietary and environmental factors and their influence on hormonal regulation during sleep deprivation, particularly emphasizing the roles of tryptophan intake and housing conditions. Overall, this research elucidates the complex relationships among sleep, dietary factors, and hormonal and inflammatory responses in rats. It underscores the potential neurotoxic effects associated with caffeine consumption when coupled with sleep deprivation. These results underscore the urgent need for further investigation into dietary strategies that could alleviate these adverse effects and promote better hormonal and overall health. By advancing our understanding of these interactions, we can pave the way for more effective interventions aimed at enhancing sleep quality and optimizing hormonal health.

## REFERENCES

AOAC (Association of Official Analytical Chemists) (2005) Official Method of Analysis of the AOAC (W. Horwitz Editor, 18^th^ Edition, Washington: D.C., AOAC).

AOAC (Association of Official Analytical Chemicals) (2006) Official Method of Analysis of the AOAC (W. Horwitz Editor, 18th Edition, Washington; D. C., AOAC).

Bellesi, M., Vivo, L. De, Chini, M., Gilli, F., Tononi, G., & Cirelli, C. (2017). Sleep loss promotes astrocytic phagocytosis and microglial activation in mouse cerebral cortex. Journal of Neuroscience, 37(21), 5263–5273. 10.1523/JNEUROSCI.3981-16.2017

Domínguez, R., Veiga-Herreros, P., Sánchez-Oliver, A. J., Montoya, J. J., Ramos-álvarez, J. J., Miguel-Tobal, F., Lago-Rodríguez, Á., & Jodra, P. (2021). Acute effects of caffeine intake on psychological responses and high-intensity exercise performance. International Journal of Environmental Research and Public Health, 18(2), 1–10. 10.3390/ijerph18020584

Donald, C. M., Moore, J., McIntyre, A., Carmody, K., & Donne, B. (n.d.). Acute Effects of 24-h Sleep Deprivation on Salivary Cortisol and Testosterone Concentrations and Testosterone to Cortisol Ratio Following Supplementation with Caffeine or Placebo. International Journal of Exercise Science, 10(1), 108–120. http://www.ncbi.nlm.nih.gov/pubmed/28479951 http://www.pubmedcentral.nih.gov/articlerender.fcgi?artid=PMC5214660

Eliot, L. (2021). Brain development and physical aggression: How a small gender difference grows into a violence problem. Current Anthropology, 62(S23), S66–S78. 10.1086/711705

Fernando, G.-P. (2008). Brain foods: the effects of nutrients on brain function. Nature Reviews Neuroscience, 9(July), 568–578. www.nature.com/reviews/neuro

Irwin, M. R., Olmstead, R., & Carroll, J. E. (2016). Sleep disturbance, sleep duration, and inflammation: A systematic review and meta-analysis of cohort studies and experimental sleep deprivation. Biological Psychiatry, 80(1), 40–52. 10.1016/j.biopsych.2015.05.014

Jenkins, T. A., Nguyen, J. C. D., Polglaze, K. E., & Bertrand, P. P. (2016). Influence of tryptophan and serotonin on mood and cognition with a possible role of the gut-brain axis. Nutrients, 8(1), 1–15. 10.3390/nu8010056

Maria, M.Y., Justo P., Julio G., Javier, V., Francisco, M. and Manuel A. (2004) Determination of tryptophan by high-performance liquid chromatography of alkaline hydrolysates with spectrophotometric detection. Food Chemistry 85(2): 317–320

Mason, G. M., Lokhandwala, S., Riggins, T., & Spencer, R. M. C. (2021). Sleep and human cognitive development. Sleep Medicine Reviews, 57, 101472. 10.1016/j.smrv.2021.101472

Moro, J., Tome, D., Schmidely, P., Demersay, T., & Azzout-Marniche, D. (2020). Histidine: A Systematic Review on Metabolism and. Nutrients, 12, 1–20.

Munteanu, I. G., & Apetrei, C. (2021). Analytical methods used in determining antioxidant activity: A review. International Journal of Molecular Sciences, 22(7). 10.3390/ijms22073380

Pan, L., An, D., & Zhu, W. Y. (2021). Sorghum as a dietary substitute for corn reduces the activities of digestive enzymes and antioxidant enzymes in pigs. Animal Feed Science and Technology, 273(January). 10.1016/j.anifeedsci.2021.114831

Park, M., Choi, Y., Choi, H., Yim, J. Y., & Roh, J. (2015). High doses of caffeine during the peripubertal period in the rat impair the growth and function of the testis. International Journal of Endocrinology, 2015, 1–9. 10.1155/2015/368475

Reuter, M., Zamoscik, V., Plieger, T., Bravo, R., Ugartemendia, L., Rodriguez, A. B., & Kirsch, P. (2021). Tryptophan-rich diet is negatively associated with depression and positively linked to social cognition. Nutrition Research, 85, 14–20. 10.1016/j.nutres.2020.10.005

Smith, I., Salazar, I., RoyChoudhury, A., & St-Onge, M. P. (2019). Sleep restriction and testosterone concentrations in young healthy males: randomized controlled studies of acute and chronic short sleep. Sleep Health, 5(6), 580–586. 10.1016/j.sleh.2019.07.003

Sousa, C., Golebiewska, A., Poovathingal, S. K., Kaoma, T., Pires-Afonso, Y., Martina, S., Coowar, D., Azuaje, F., Skupin, A., Balling, R., Biber, K., Niclou, S. P., & Michelucci, A. (2018). Single-cell transcriptomics reveals distinct inflammation-induced microglia signatures. EMBO Reports, 19(11), 1–17. 10.15252/embr.201846171

Spiegel, K., Knutson, K., Leproult, R., Tasali, E., & Van Cauter, E. (2005). Sleep loss: A novel risk factor for insulin resistance and Type 2 diabetes. Journal of Applied Physiology, 99(5), 2008–2019. 10.1152/japplphysiol.00660.2005

Tardy, A. L., Pouteau, E., Marquez, D., Yilmaz, C., & Scholey, A. (2020). Vitamins and minerals for energy, fatigue and cognition: A narrative review of the biochemical and clinical evidence. Nutrients, 12(1). 10.3390/nu12010228

Van Cauter, E., Spiegel, K., Tasali, E., & Leproult, R. (2008). Metabolic consequences of sleep and sleep loss Sleep Medicine 2008 Vol 9. Sleep Med, 9(suppl 1), S23–S28.

Wan, Y., Gao, W., Zhou, K., Liu, X., Jiang, W., Xue, R., & Wu, W. (2022). Role of IGF-1 in neuroinflammation and cognition deficits induced by sleep deprivation. Neuroscience Letters, 776(March), 136575. 10.1016/j.neulet.2022.136575

